# Genetic stability of *Aedes aegypti* populations following invasion by *w*Mel *Wolbachia*

**DOI:** 10.1101/2021.06.03.446908

**Authors:** Meng-Jia Lau, Tom Schmidt, Qiong Yang, Jessica Chung, Lucien Sankey, Perran A. Ross, Ary A. Hoffmann

## Abstract

**Background:** *Wolbachia w*Mel is the most used strain in mosquito rear and release strategies that aim to inhibit the transmission of arboviruses such as dengue, Zika, Chikungunya and yellow fever. However, the long-term establishment of *w*Mel in natural populations of the dengue mosquito *Aedes aegypti* raises concerns that interactions between *Wolbachia w*Mel and *Ae. aegypti* may lead to changes in the host genome, which could affect useful attributes of *Wolbachia* that allow it to invade and suppress disease transmission.

**Results:** We applied an evolve-and-resequence approach to study genome-wide genetic changes in *Ae. aegypti* from the Cairns region, Australia, where *Wolbachia w*Mel was first introduced more than 10 years ago. Mosquito samples were collected at three different time points in Gordonvale, Australia, covering the phase before (2010) and after (2013 and 2018) *Wolbachia* releases. An additional three locations where *Wolbachia* replacement happened at different times across the last decade were also sampled in 2018. We found that the genomes of mosquito populations mostly remained stable after *Wolbachia* release, with population differences tending to reflect the geographic location of the populations rather than *Wolbachia* infection status. However, outlier analysis suggests that *Wolbachia* may have had an influence on some genes related to immune response, development, recognition and behavior.

**Conclusions:** *Aedes aegypti* populations remained geographically distinct after *Wolbachia* releases in North Australia despite their *Wolbachia* infection status. At some specific genomic loci, we found signs of selection associated with *Wolbachia*, suggesting potential evolutionary impacts can happen in the future and further monitoring is warranted.

## Background

*Wolbachia* are bacteria that live inside the cells of many insects and induce a number of important phenotypic effects on their hosts that can be harnessed for pest and disease control. *Wolbachia*-infected *Aedes aegypti* mosquitoes have now been released in multiple locations of the world [1–3] to help reduce the transmission of arboviruses such as dengue, Zika, Chikungunya and yellow fever [4–6]. *Wolbachia w*Mel, which was transferred artificially from *Drosophila melanogaster* into *Ae. aegypti* [6], was first released in Gordonvale and Yorkeys Knob, Queensland, Australia, around a decade ago [7] where it invaded the local population through cytoplasmic incompatibility (CI). CI results in uninfected females less likely to produce viable offspring if they mate with infected males. In contrast, infected females produce viable offspring when they mate with uninfected males or males infected by the same *Wolbachia* strain, and these offspring are infected [8]. This allows *Wolbachia* to invade and be self-sustained in a population but may also increase population divergence because it can reduce the “effective migration rate” [9, 10] between infected and uninfected populations. *Wolbachia* can also impact mitochondrial DNA (mtDNA) variation through indirect linkage disequilibrium [11–13].

With the success of *Wolbachia* in suppressing dengue following invasion [1, 3], many studies have now focused on the sustainability of this approach beyond the initial spread, such as the maintenance of high infection levels [14], fitness costs [15, 16] and evolutionary adaptation [17, 18]. The potential evolutionary changes in *Wolbachia w*Mel-infected *Ae. aegypti* as well as in the bacterial genome itself following releases in the field have raised concerns about the long-term effectiveness of the strategy. The genetic background of the mosquito host can affect the capacity of *Wolbachia* to invade populations and suppress arboviruses [2, 19, 20]. *Aedes aegypti* has a short generation interval of ~1 month, and so if the introduction of *Wolbachia* triggers an evolutionary process in the mosquito genome this could be observable within a few years after invasion. Evolutionary changes in response to natural *Wolbachia* infections have previously been noted and appear to involve both the *Wolbachia* and host genomes [21, 22], affecting the population dynamics of *Wolbachia* infections. Adaptations can occur to counter any negative fitness effects of *Wolbachia*, as documented in *Drosophila* [21, 23], and negative fitness effects are particularly evident in novel infections transfected into new hosts [24].

In the past decade, the *w*Mel infection itself has not evolved in terms of either sequence composition or density since establishment in *Ae. aegypti* in northern Queensland, Australia [25]. Phenotypic comparisons also suggest limited changes in host fitness costs and CI since population replacement in this region [14, 18], although the number of fitness-related traits scored so far has been limited. Blockage of virus transmission also appears stable to date [26, 27], and may persist through ongoing selection favoring high viral blocking in *Ae. aegypti* populations [19].

There are no published studies investigating evolutionary changes in wild host populations at the genomic level following a *Wolbachia* release. Any putative changes may guide further phenotypic comparisons based on the types of candidate genes identified. However, there are challenges in characterizing genome changes in *Ae. aegypti* after *Wolbachia w*Mel infection. First, the genome of *Ae. aegypti* contains a large proportion (47%) of transposable elements (TEs), which result in a large genome size (1.38Gb) compared to other mosquitoes [28–30]. TEs might also enhance rates of evolution, given that they are involved with gene regulation, and increase genome plasticity [31]. Moreover, the involvement of other environmental factors in field-collected samples cannot be neglected, impacts of gene flow following the activity of *Wolbachia* release will also increase the difficulty of outlier analyses. Finally, compared to model organisms, the genome of *Ae. aegypti* is still relatively poorly annotated, with only 256 proteins (<1%) reviewed in the Swiss-Prot database (https://www.uniprot.org/taxonomy/7159).

In this study, we analyse pooled whole genome sequencing (WGS) data of mosquitoes from Gordonvale, Australia, where releases first took place, covering three different time points from the pre- and post-release phase. As a comparison, we also sequenced samples in Edge Hill, Redlynch and Yorkeys Knob, where *Wolbachia* replacement happened at different times across the last decade. We combined analyses of spatial variation in *Wolbachia* infection status and of genetic diversity in the mosquitoes to reveal the potential impact of *Wolbachia w*Mel on the genome of its host *Ae. aegypti*.

## Results

### Genetic variation in *Aedes aegypti* populations

*Aedes aegypti* were collected from four sites around Cairns, Australia, at different times pre- and post-*Wolbachia* release (Fig. 1, Table 1). We investigated patterns of genetic variation within populations using PoPoolation v. 1.2.2 [32] with the genomic annotation file from the reference AaegL5.0. In Tajima’s pi (nucleotide diversity *π*) analysis (Fig. 1), differences we found among the populations depended on chromosomal location. We analysed the genome-wide nucleotide diversity in 10 kbp non-overlapping windows. The nucleotide diversities were different among populations, while in areas near the center of each chromosome, diversities were low and similar for all populations. This pattern has been shown in previous sequencing studies [30, 33]. Overall, nucleotide diversity was highest on chromosome 1, which contains the sex determining locus and contains relatively lower gene densities and TEs but higher satellites compared to chromosomes 2 and 3 [34].

**Fig. 1.**
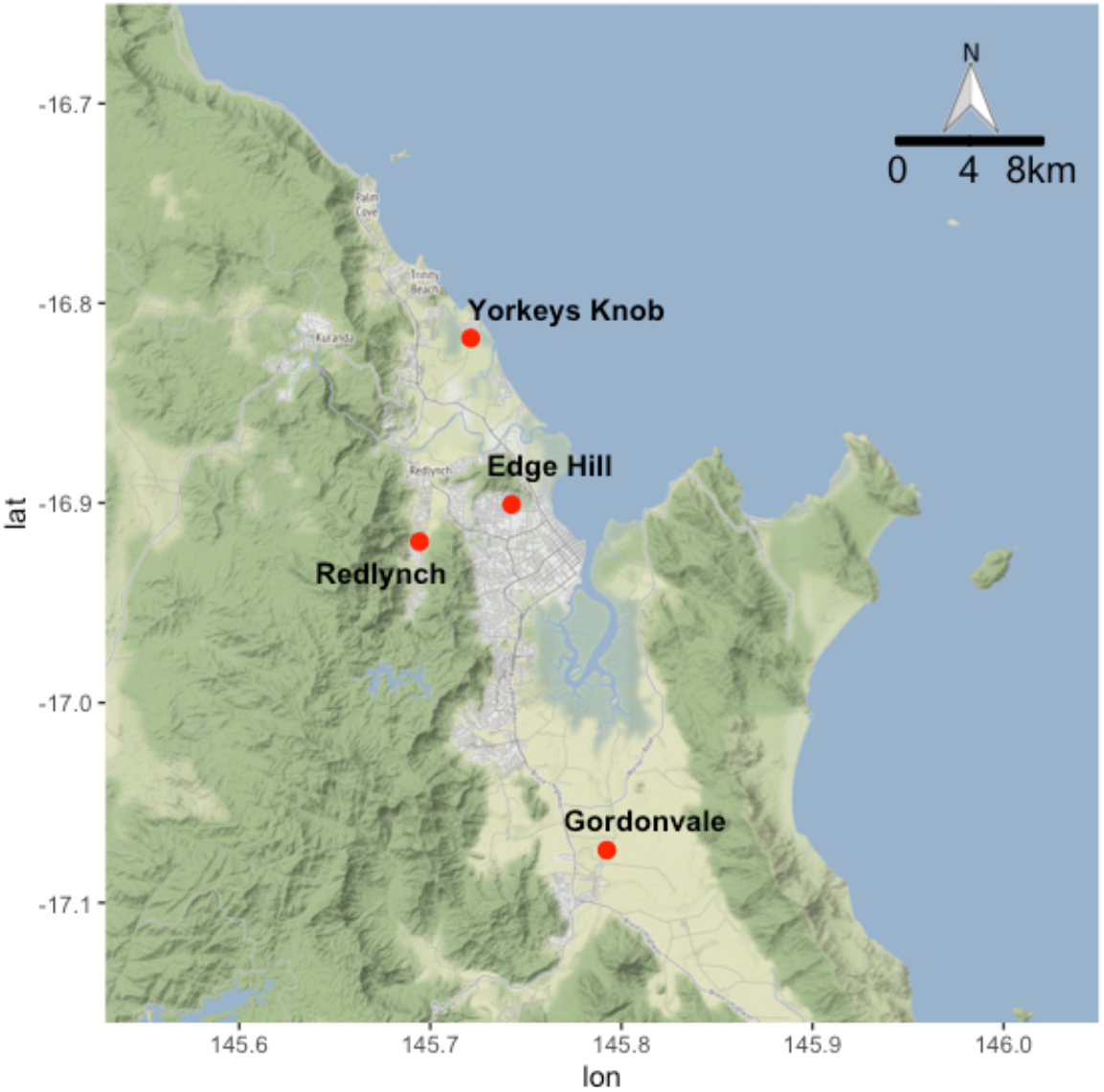
Locations of sampled *Aedes aegypti* populations. Samples from Gordonvale were collected in 2010, 2013 and 2018, while samples from other locations were collected in 2018. Axes show longitude (lon) and latitude (lat).

**Table 1.**
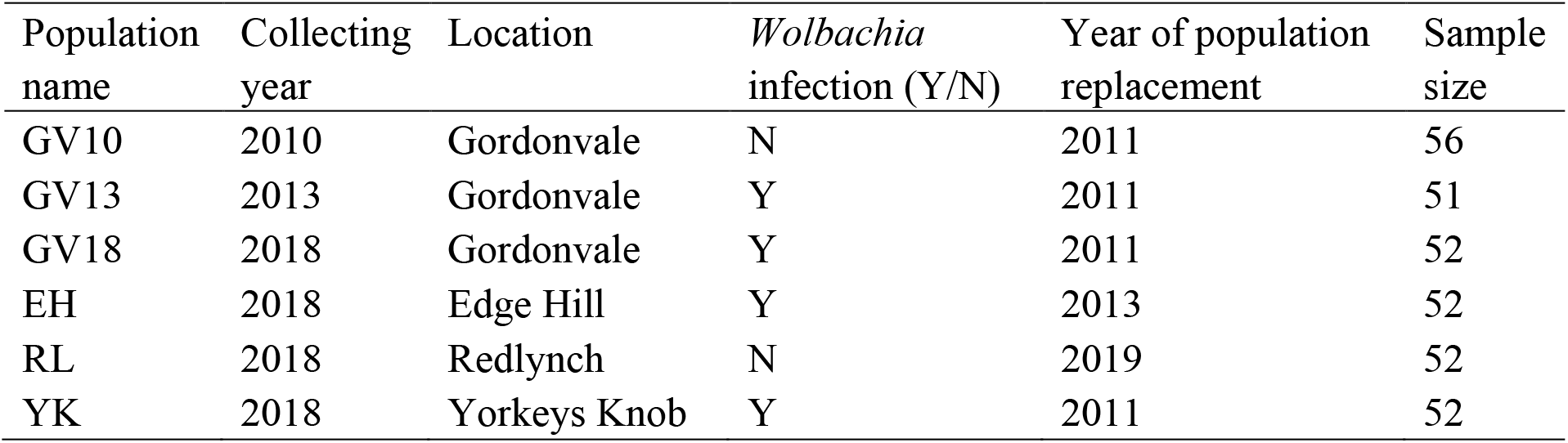
Summary of *Aedes aegypti* collections and designations of samples used in comparisons.

| Population name | Collecting year | Location | *Wolbachia* infection (Y/N) | Year of population replacement | Sample size |
| --- | --- | --- | --- | --- | --- |
| GV10 | 2010 | Gordonvale | N | 2011 | 56 |
| GV13 | 2013 | Gordonvale | Y | 2011 | 51 |
| GV18 | 2018 | Gordonvale | Y | 2011 | 52 |
| EH | 2018 | Edge Hill | Y | 2013 | 52 |
| RL | 2018 | Redlynch | N | 2019 | 52 |
| YK | 2018 | Yorkeys Knob | Y | 2011 | 52 |

In Tajima’s D analysis, we found that the density distributions of values were similar between populations, except for Gordonvale pre-release (GV10) which shows a high proportion of negative values (Fig. 3). This pattern is more obvious at the gene level (Fig. 3b) than at the genome level measured in 10 kbp non-overlapping windows (Fig. 3a). The four 2018 populations converge regardless of their *Wolbachia* status or time since *Wolbachia* was invaded. This suggests that the pattern reflects a difference in GV10 before release rather than an effect of *Wolbachia per se*.

**Fig. 2.**
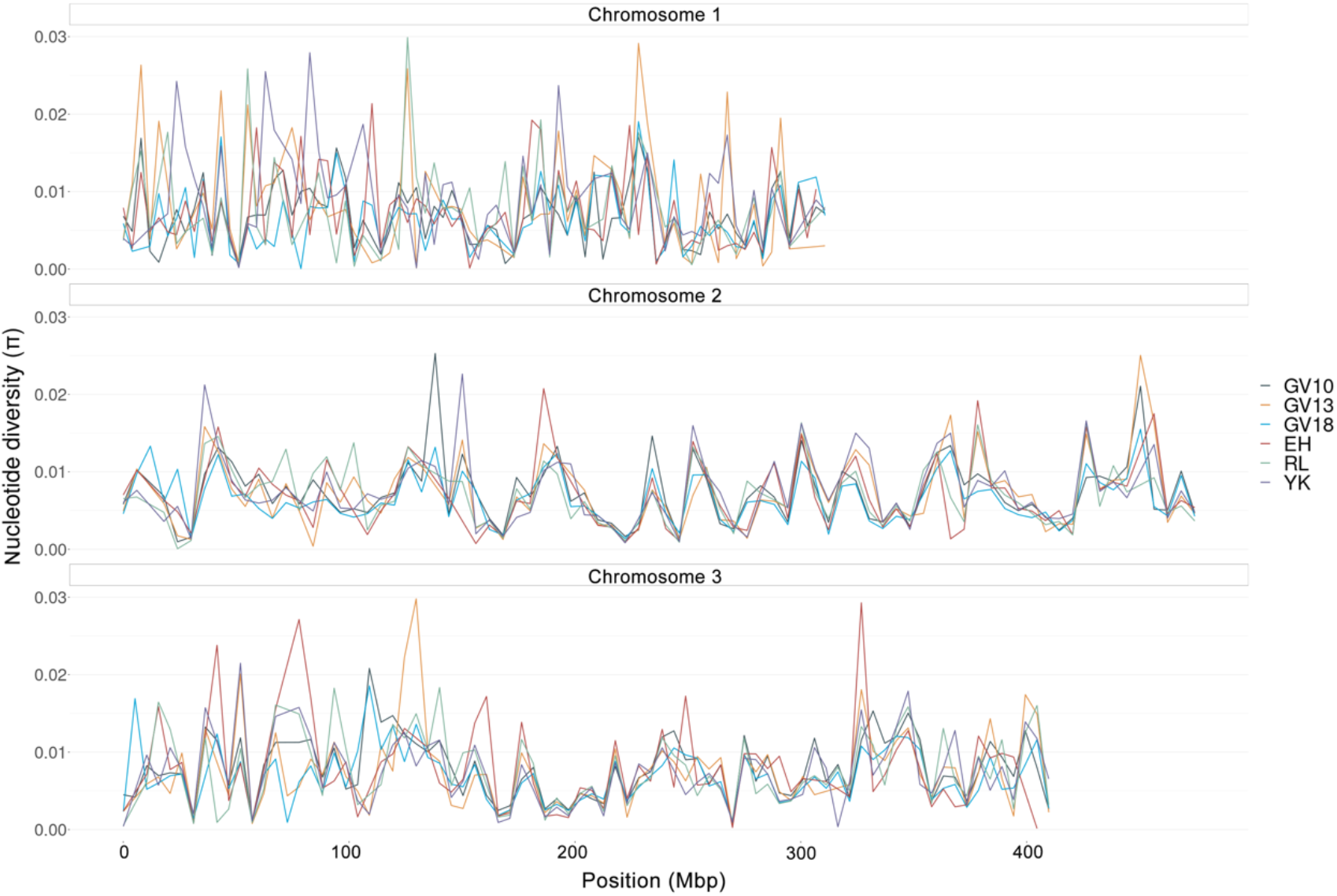
LOESS-smoothed curves of genome-wide nucleotide diversity (*π*). Six populations of *Ae. aegypti* measured in 10 kbp non-overlapping windows. GV10 and GV13 represent samples collected in 2010 and 2013 from Gordonvale; GV18, EH, YK and RL represent samples collected in 2018 from Gordonvale, Yorkeys Knob, Edge Hill and Redlynch respectively.

**Fig. 3.**
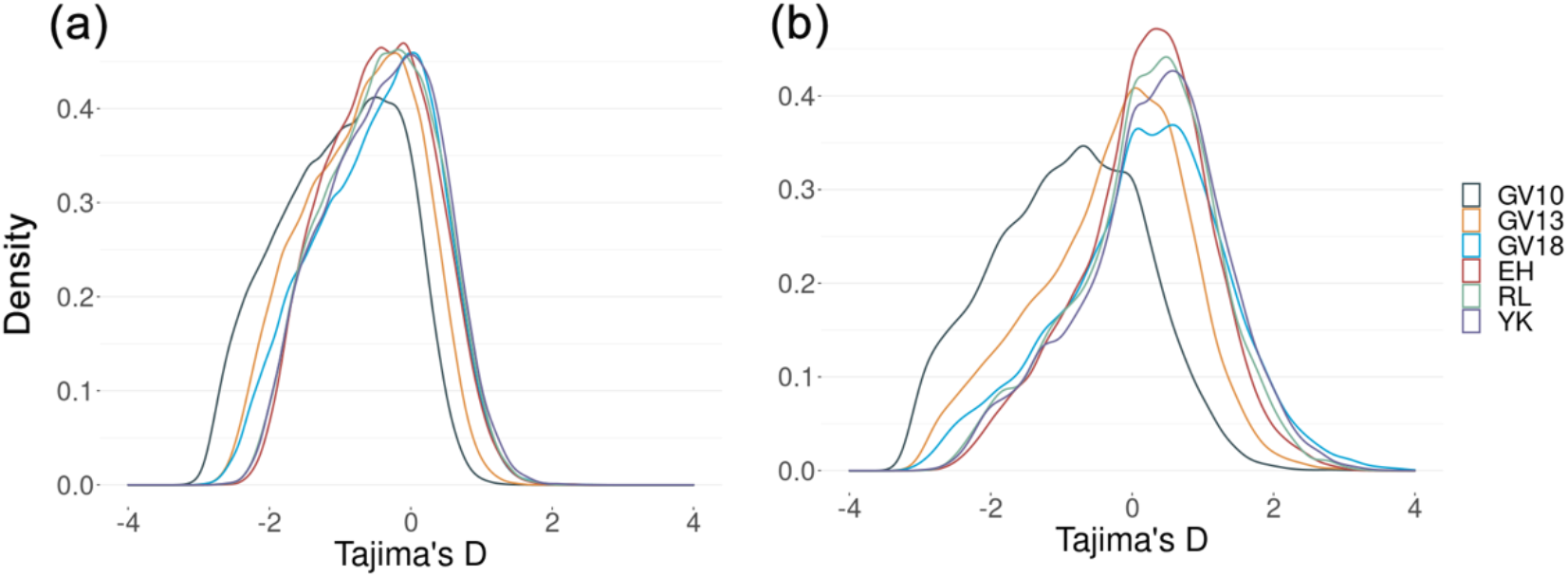
Density distributions of Tajima’s D values. Tajima’s D measured at (a) the genome level measured in 10 kbp non-overlapping windows and at (b) the gene level. GV10 and GV13 represent samples collected in 2010 and 2013 from Gordonvale; GV18, EH, YK and RL represent samples collected in 2018 from Gordonvale, Yorkeys Knob, Edge Hill and Redlynch respectively.

### Geographic segregation of *Aedes aegypti* populations

We obtained a genetic distance matrix from the average of pairwise Fst (Fixation index) values through 100 kbp non-overlapping windows (Additional file 1), with temporally-separated GV samples tending to have lower Fst values than comparisons across geographically separated samples. In the analysis of isolation by distance, with only four 2018 populations included, the genetic distance had a weak but nonsignificant correlation with log transformed geographical distance (road distance) in a Mantel test (r=0.66, p=0.12).

There were 461,067 SNPs left after filtering with minimal depth of 50 in all populations and an average MAF (minor allele frequency) more than 0.1. We did a principal components analysis based on these SNPs, with PC1 and PC2 accounting for 24.2% and 22.9% of the variance respectively (Fig. 4a). The three temporally-seperated GV samples fell out together, but GV10 was closer to GV18 than to GV13. We also did a PCA analysis on pairwise Fst differences in 100 kbp non-overlapping windows across the genome (Fig. 4b, d) or at the gene level (Fig. 4c, e). When testing the similarity of infected or uninfected populations (Fig. 4d, e), we found little evidence for any clustering of populations related to *Wolbachia* infection status either across the genome or at the gene level. On the other hand, these populations were separated geographically (Fig. 4b, c); the pairwise comparisons of GV samples and samples located away from GV tended to cluster when comparing across the genome, and were more separated at the gene level.

**Fig. 4.**
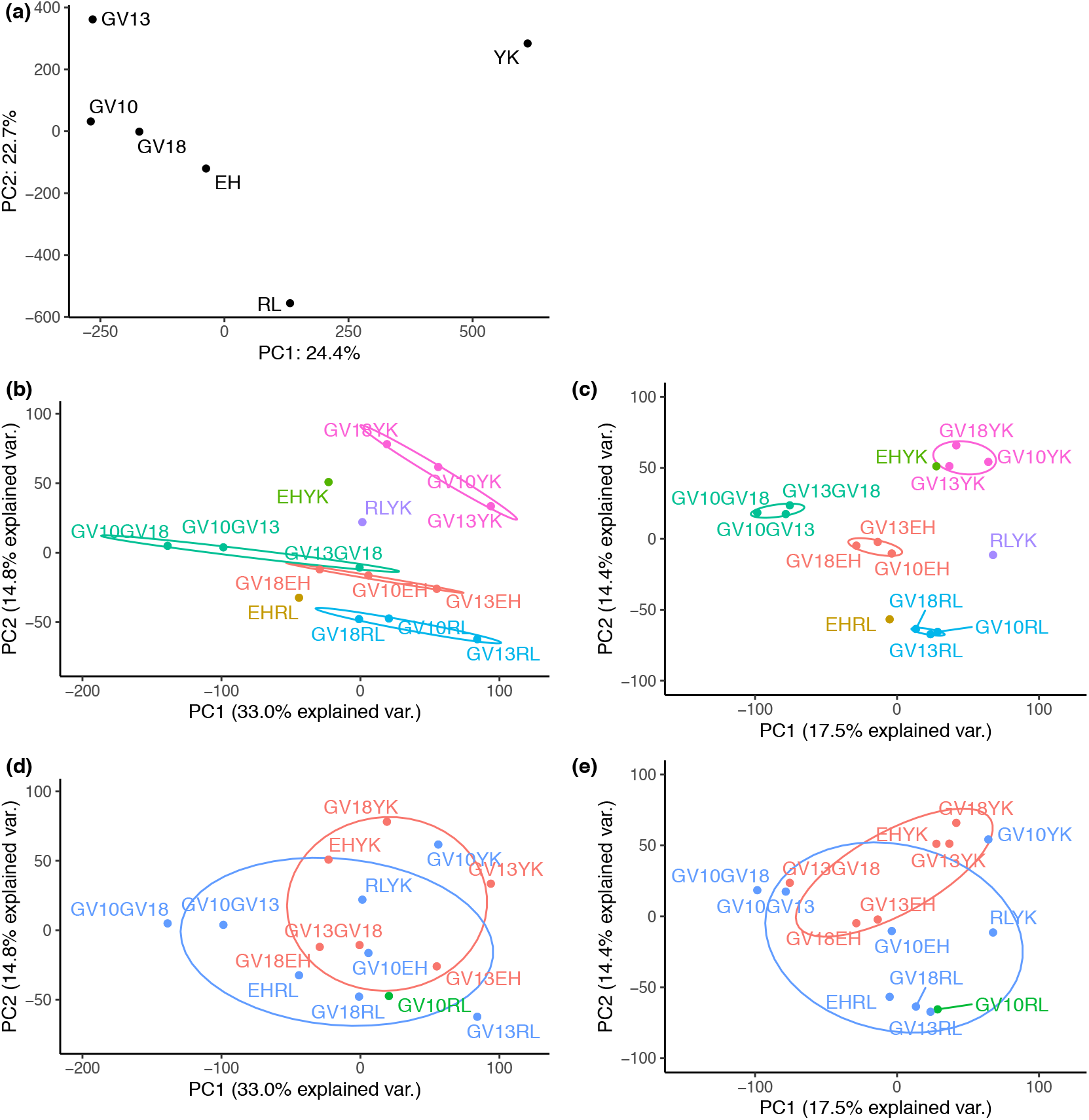
Principal components analysis based on MAF or pairwise Fst. PCA plots of (a) allele frequency of *Aedes aegypti* samples (MAF > 0.1, minimal coverage > 50); (b, d) pairwise Fst throughout the genome with 100 kbp non-overlapping windows, and (c, e) pairwise Fst within genes. Colors in (b, c) represent comparisons between GV samples and a sample from a different location. The blue color in (d, e) represents comparisons between *Wolbachia*-infected and uninfected samples, while red represents comparisons within *Wolbachia*-infected samples and green represents the comparison within uninfected samples.

### Bayesian models to identify outliers potentially associated with *Wolbachia*

To investigate potential selection associated with *Wolbachia*, we introduced 461,067 SNPs from the above filtering process and used two Bayesian models from BayPass v. 2.2 [35] for *Wolbachia*-related outlier analysis.

In the covariate model of BayPass, with *Wolbachia* infection status in the comparison, we found 2415 SNPs showing a “substantial” signature of selection with an average BF * > 5 (Bayes Factor in dB units (BF*= 10 × log10(BF)), and 391 showing “strongly-selected” signature of selection with an average BF* > 10 (Fig. 5).

**Fig. 5.**
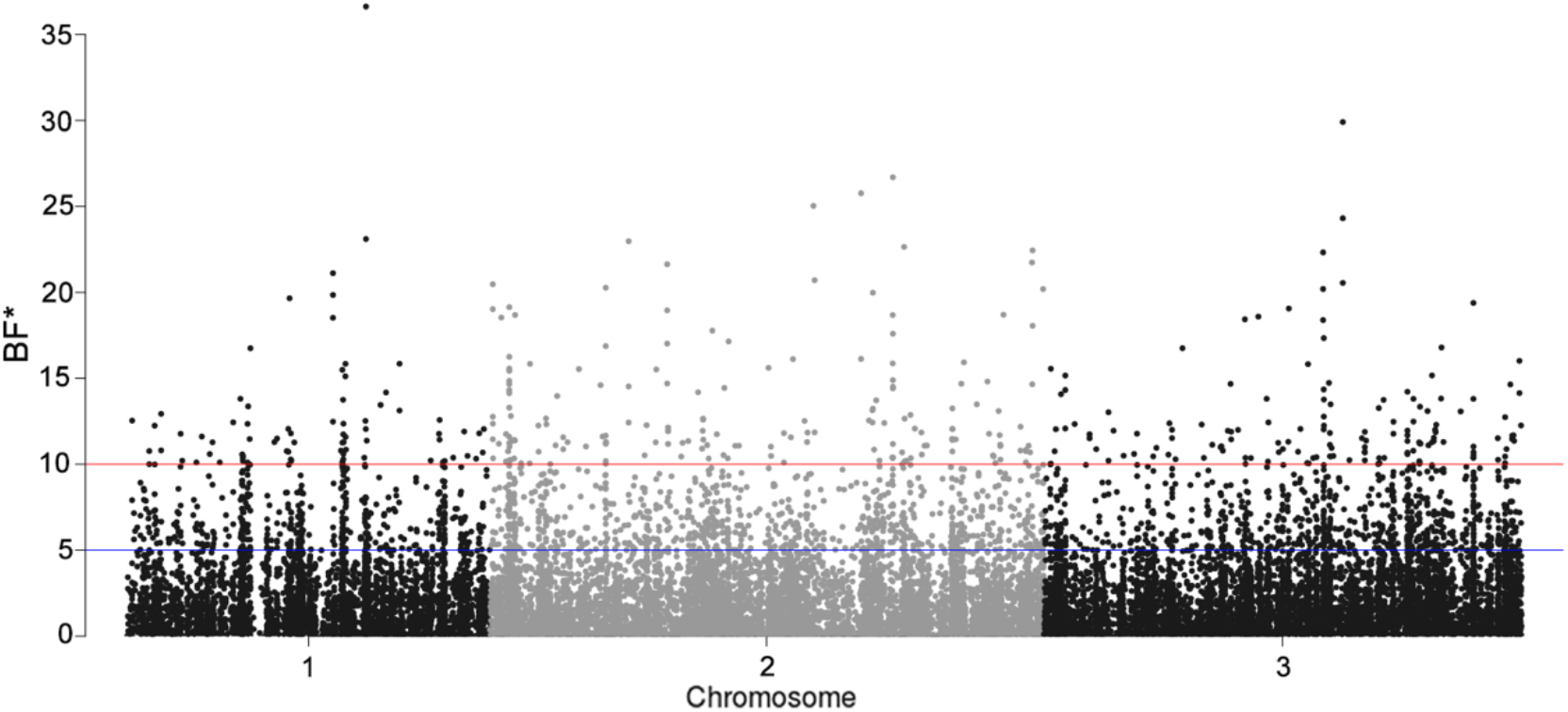
Manhattan plot of SNPs in covariate model representing *Wolbachia* infection status. Only SNPs with a positive Bayes Factor are shown. The horizontal blue lines represent the cut-off for SNPs with BF* > 5 (2415 outliers) and the horizontal red lines represent the cut-off for SNPs with BF* > 10 (391 outliers).

The introduction of linear relationships in the covariate model, however, can have high false positives from sampling noise in particular when large environmental effects are involved and small number of populations are compared. We therefore used a second model to calculate X^T^X, which was analogous to Fst [35, 36], then combined these two models to identify outliers potentially associated with *Wolbachia* infection.

We found that 950 (prior probability: 0.74 in each Mbp) of the “substantial” SNPs fell into the intersection of top 10% X^T^X values in the comparisons between GV10 and GV13 and between GV10 and GV18, including 229 SNPs distributed on chromosome 1, 390 on chromosome 2 and 330 on chromosome 3, while one was found on an assembly scaffold NW_018735222.1. A proportion of these SNPs were concentrated in some regions, with posterior probability at least five times greater than prior probability (> 3.7 in each Mbp), suggesting selection on *Wolbachia* (Table 2). These SNPs were considered as “substantial” outliers associated with the *Wolbachia* infection.

**Table 2.** Selected regions impacted by *Wolbachia* infection status.

| Chromosome | Position | Size (Mbp) | Number of SNPs |
| --- | --- | --- | --- |
| 1 | 96900000-105500000 | 8.6 | 39 |
| 1 | 138000000-140500000 | 2.5 | 13 |
| 1 | 148500000-149500000 | 1.0 | 11 |
| 1 | 183000000-188300000 | 5.3 | 40 |
| 1 | 203100000-206200000 | 3.1 | 22 |
| 2 | 15600000-16800000 | 1.2 | 27 |
| 2 | 20700000-21200000 | 0.5 | 11 |
| 2 | 98800000-99300000 | 0.5 | 18 |
| 2 | 345200000-345400000 | 0.2 | 19 |
| 2 | 396100000-396500000 | 0.4 | 15 |
| 3 | 239700000-249400000 | 9.7 | 31 |
| 3 | 311900000-316000000 | 4.1 | 25 |
| 3 | 368400000-368600000 | 0.2 | 14 |

We also used a stricter criterion: average BF*>=10 and X^T^X values falling into the intersection of 95% threshold in the GV10, GV13 comparison and the GV10, GV18 comparison. We then found 113 SNPs that were highly associated with *Wolbachia* infection and were considered as “strong” outliers.

### Pathway analysis and gene ontology enrichment analysis

The “substantial” outliers were distributed within 187 genes (Additional file 2). There were 1436 genes potentially involved when a 100-kbp region around the outliers was taken into account (Additional file 3). We performed a pathway analysis of these 1436 genes through the KEGG database and found eight pathways significantly involved (Table 3, Additional file 4), including pathways involved with development and immune response (MAPK signalling pathway and Toll and Imd signaling pathway). These genes were also BLAST searched for homologues in *Drosophila melanogaster*; after filtering out the low-quality matches, we obtained 577 homologues proteins (Additional file 5). In the gene ontology enrichment (GO) analysis, 108 gene sets were significantly enriched in biological process (Additional file 6), 42 in cellular components (Additional file 7) and 22 in molecular factors (Additional file 8). In general, we found that *Wolbachia w*Mel may modulate genes with diverse functions such as cell development, interaction, cellular response, cellular transport, neurogenesis, lipid and glucide metabolization, behavior and immune response.

**Table 3.** Significantly enriched pathways in KEGG database potentially involved with *Wolbachia* infection.

| Pathway | Description | Gene Ratio | Bg Ratio | P value |
| --- | --- | --- | --- | --- |
| aag00232 | Caffeine metabolism | 6/296 | 12/3281 | <0.001 |
| aag00981 | Insect hormone biosynthesis | 9/296 | 40/3281 | 0.008 |
| aag00052 | Galactose metabolism | 7/296 | 30/3281 | 0.015 |
| aag04341 | Hedgehog signaling pathway - fly | 6/296 | 25/3281 | 0.021 |
| aag04013 | MAPK signaling pathway - fly | 14/296 | 92/3281 | 0.034 |
| aag00230 | Purine metabolism | 14/296 | 96/3281 | 0.047 |
| aag00270 | Cysteine and methionine metabolism | 6/296 | 30/3281 | 0.048 |
| aag04624 | Toll and Imd signaling pathway | 9/296 | 54/3281 | 0.050 |

The “strong” outliers were distributed across 31 genes (Table 4), including some interesting ones, such as cytochrome P450 gene 5564751, which was found enriched in response to dengue virus infection in refractory mosquitoes [37], and associated with insecticide resistance [38, 39]. Carbonic anhydrase gene 5565700 has the function of balancing pH in mosquito midgut [40, 41]. Ecdysone protein E75, encoded by gene 5569135, and lipophorin, encoded by gene 5572681, were highly expressed in females after blood feeding, potentially involved in regulation of oogenesis and vitellogenesis [42, 43]. Charged multivesicular body protein, encoded by gene 5573292, was associated with endosome formation and can influence mosquito immune response [44].

**Table 4.**
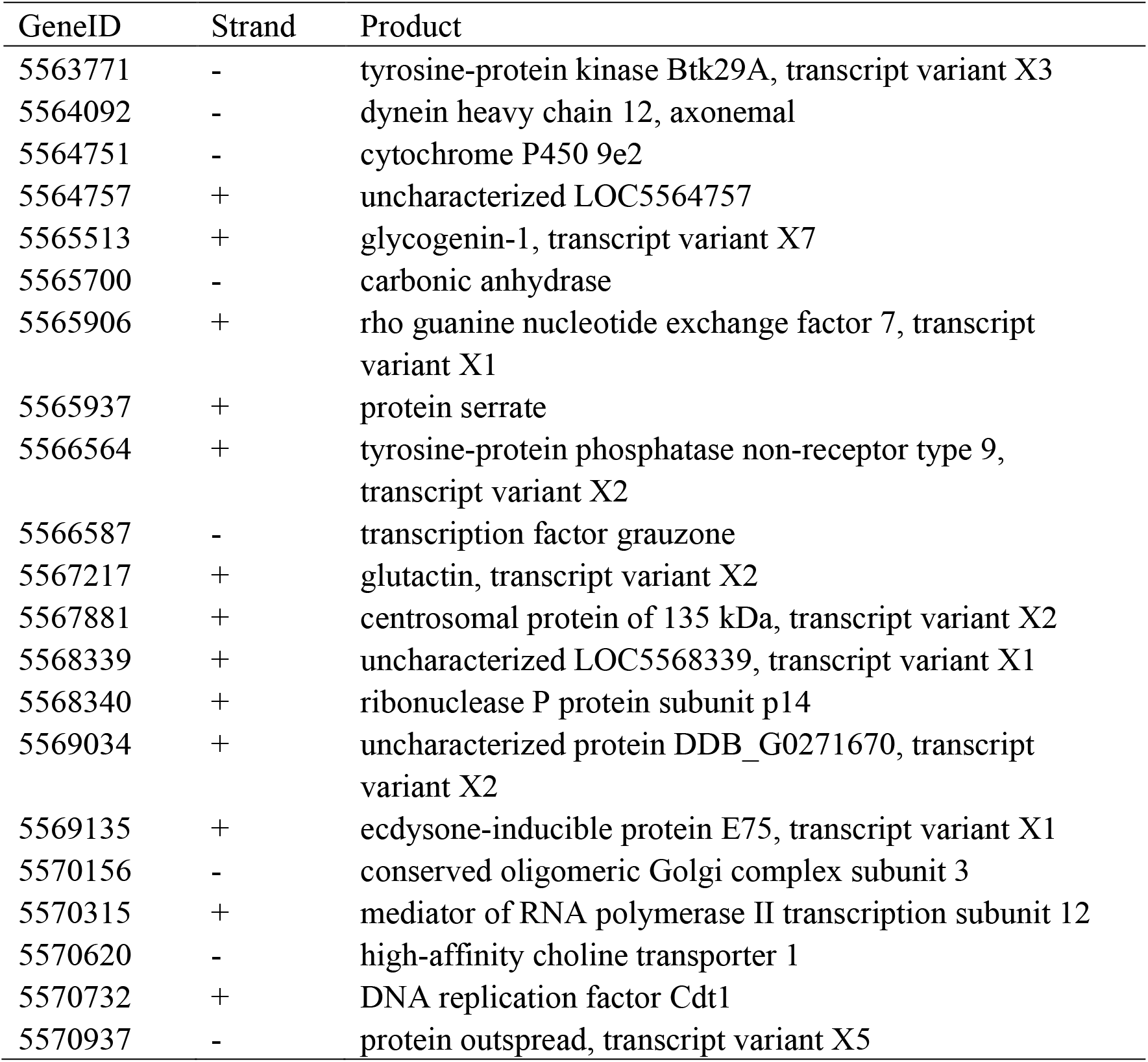
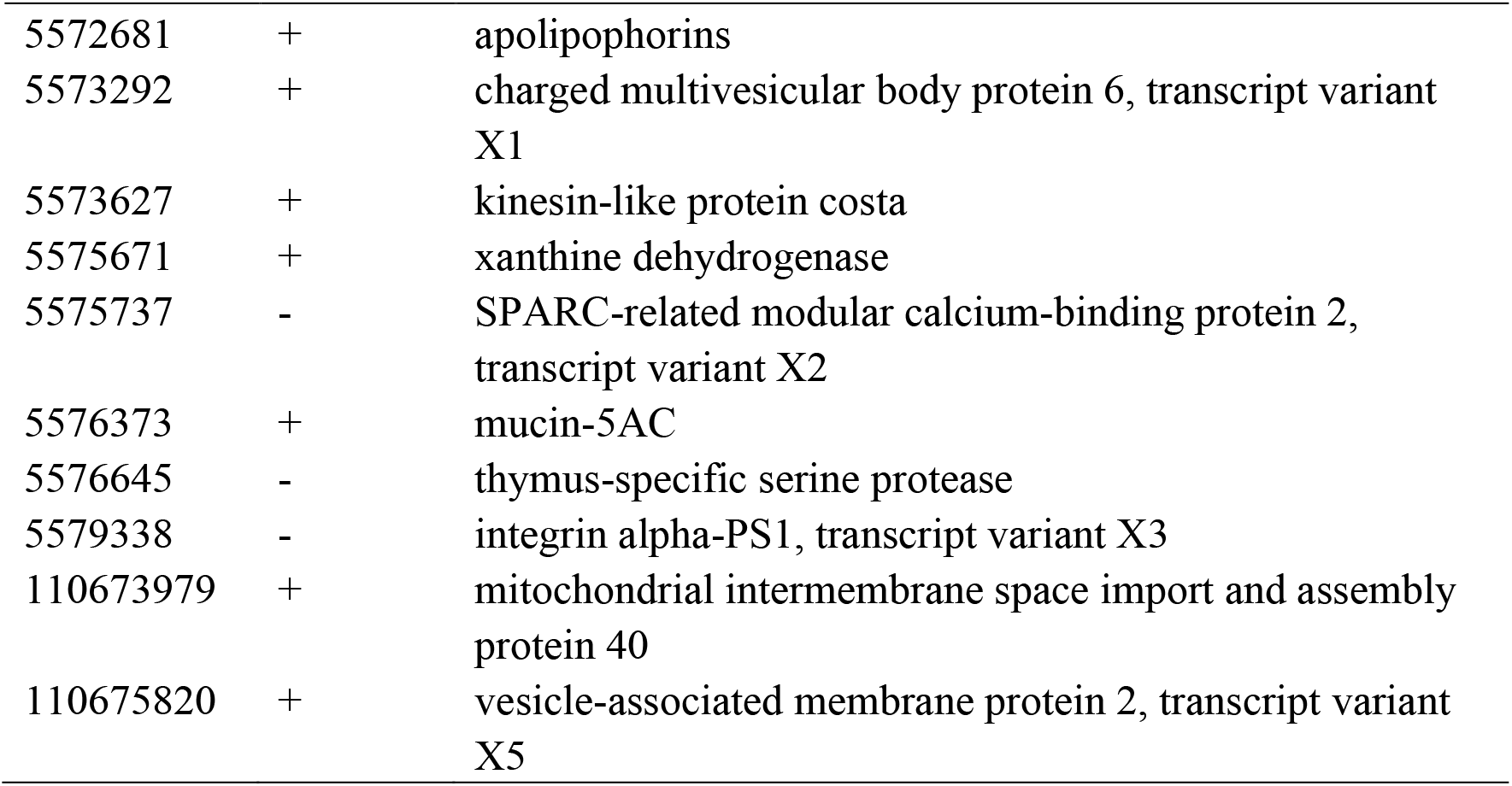
“strong” outliers associated with *Wolbachia* infection are distributed in 31 genes.

## Discussion

We show that *Ae. aegypti* populations in Cairns remain geographically distinct following releases of *w*Mel, but also find some suggestive evidence for evolutionary changes in mosquito populations. When interpreting the results, it is important to consider the release process and target population and the fact that *Wolbachia*-induced CI can increase population divergence by reducing the migration rate across host populations when only one or both (in the case of different *Wolbachia* strains) are infected [9, 10]. The population released involved a *w*Mel transinfected strain that had been repeatedly backcrossed to a Cairns field population background, with the expectation that the released background would be 99.9% Cairns [7]. For releases in Yorkeys Knob and sites around Cairns, we did not expect the release population to differ from the background population because there is movement of mosquitoes around this area as reflected by the occasional detection of *Wolbachia*-infected mosquitoes after the release [7]. On the other hand, Gordonvale is a relatively isolated population which may have its own seasonal dynamics. Although this population is not genetically isolated based on microsatellite and EPIC markers [45], it does appear to be somewhat separated based on the SNP markers used in the current study. This may account for the pattern noted for Tajima’s D where the 2010 Gordonvale population was a clear outlier.

When releasing mosquitoes and following invasion by *Wolbachia*, there is not only complete replacement of the uninfected mosquito population by *Wolbachia*-infected mosquitoes but also replacement of the mtDNA that can hitchhike along with the *Wolbachia* [12]. Also, while any linkage disequilibrium between the *Wolbachia* and nuclear DNA variants is expected to break down relatively quickly [46], new alleles may nevertheless be introduced into the population. The nuclear DNA constitution of the population might be expected to become more like the release stock for a period as released females and their offspring mate with released and resident males, although local selection should then lead to populations becoming more like the original population. In our case, the genetic similarity between the Gordonvale populations after release and the original population prior to release might reflect local selection and ongoing introgression of the release stock with the resident population, as GV10 and GV18 are closer than GV13 in the PCA analysis. Furthermore, Yorkeys Knob and Edge Hill remain distinct from each other despite previously being invaded by the same release stock [3, 7].

Populations were more segregated at the gene level than at the genome level, which may be a consequence of a high mobility of TEs associated with ORF regions [21]. On the other hand, selection in response to local conditions and the impact of *w*Mel on *Ae. aegypti* may nevertheless influence patterns of genetic differentiation at specific loci [19]. Our Bayesian outlier analysis identified several regions in each chromosome and genes related to immune response, development, recognition and behavior that may have been under selection. These potential evolutionary impacts of *Wolbachia w*Mel on the genome of *Ae. aegypti* in the field suggest that further monitoring is warranted, although at this stage other factors unrelated to *Wolbachia* appear to have a larger impact on genomic differentiation among samples.

We found signs of selection on the Toll and Imd signalling pathway in KEGG analysis; these are important pathways in immune systems [47, 48] and viral blocking processes [48–50]. Previous transcriptomic studies of *Wolbachia* in *Ae. aegypti* showed up-regulation of these pathways in both *w*Mel and *w*MelPop-infected *Ae. aegypti* [51, 52]. Viral blocking genes are mainly distributed on chromosome 1, in addition to genes related to cytoskeleton, cell-cell adhesion and signal transduction [19]; these genes also show up in our GO analysis. Other than viral blockage, caffeine metabolism was strongly impacted, which may impact hormone metabolism and detoxification when cytochrome P450 is involved [53, 54]. We also detected enriched pathways involved with development, such as insect hormone biosynthesis and the Hedgehog signaling pathways. In GO enrichment analysis these are represented in cell growth, structure, recognition and behavior.

## Conclusions

*Wolbachia w*Mel-infected *Ae. aegypti* mosquitoes have been released successfully in the field to help reduce the transmission of arboviruses, but interactions between *w*Mel and *Ae. aegypti* could result in adaptation [55, 56], altering viral blocking efficiency [19, 57], host fecundity [21] and insecticide resistance [58]. In this study, we have identified *Ae. aegypti* populations as being geographically distinct despite their *Wolbachia* infection status. However, selection associated with *Wolbachia* may still have influenced variation at some loci. This is the first time that genome evolution associated with *Wolbachia* infection has been examined in field populations where there has been a deliberate release. However, it is hard to draw conclusions about long-term impacts of *Wolbachia* on the *Aedes* genome, which may take more time to develop, and which may be different in regions where dengue is endemic, unlike in Australia. Our findings highlight the possibility that the effect of *Wolbachia* can interact with the host genomic background, which has been shown previously based on phenotypic assays on the longevity of *w*MelPop in *Drosophila* [59].

## Methods

### Samples and study sites

*Aedes aegypti* samples were collected from four sites around Cairns (Fig. 1, Table 1). In Gordonvale, samples were collected three times: in the summer of 2010 (prerelease), as well as in 2013 and 2018 (2 – 7 years post release given that the area was stably invaded in 2011 [7]). Samples from Yorkeys Knob, Edge Hill and Redlynch were collected in 2018. Yorkeys Knob experienced *Wolbachia* invasion at the same time as Gordonvale in 2011, and Edge Hill which was invaded in 2013 [60]; Redlynch was an uninfected area when sampled in 2018. Gordonvale samples from 2010 and 2013 were collected by BG-Sentinel traps (Biogents, Regensburg, Germany) while 2018 samples were collected by ovitraps, taking care to sample only 1-2 larvae per ovitrap to reduce the likelihood of siblings being sampled [61]. Samples from Gordonvale 2010 (GV10) and 2013 (GV13) were stored in 100% ethanol at the adult stage while samples collected from 2018 (GV18, EH, YK and RL) were stored at the fourth instar larval stage.

### DNA extraction and library preparation

Whole genomic DNA was extracted from each individual mosquito using Qiagen DNA Blood and Tissue kit (Venlo, Limburg, NL) for 2010 and 2013 samples, using Roche High Pure™ PCR Template Preparation Kits (Roche Molecular Systems, Inc., Pleasanton, CA, USA) for 2018 samples. *Wolbachia* infection status was confirmed by a diagnostic qPCR test as outlined elsewhere [62]. The concentration of extracted individual DNA was measured using Quantitation Qubit™ 1X dsDNA HS Assay Kit (Invitrogen Life Technologies USA). Samples from each of the six populations (Table 4) were pooled prior to sequencing based on an equal amount of DNA from each individual. Each population was sent for whole genome sequencing with > 50 depth via Illumina Hiseq2500 using 100 bp paired read chemistry for GV10 and GV13, and 150 bp paired read chemistry for GV18, EH, YK and RL libraries.

Raw sequences were trimmed using Trimmomatic v. 0.39 to truncate and remove low quality reads; the reads with a phred score above 20 and length above 70 were kept. The reference genome AaegL5.0 [30] was indexed and reads were aligned to the reference by bowtie2 v. 2.3.4.3 with the very-sensitive-local mode [63]. Samtools v.1.9 was used with default parameters to sort, mark and generate pileup files requiring a minimal mapping quality of 20.

### Estimation of genome variation

We investigated patterns of genetic variation within populations using PoPoolation v. 1.2.2 [32] with the genomic annotation file from the reference AaegL5.0. We calculated Tajima’s pi (nucleotide diversity π) for each population at 10 kbp non-overlapping windows when the coverage was not less than 20. Windows with low coverage generated no values and were neglected before adjusting the shape of lines across each chromosome by a LOESS smooth curve [64]. We also calculated Tajima’s D for each population at 10 kbp non-overlapping windows and at the gene level with a minimal coverage of 20. The value of Tajima’s D was calculated from allele frequencies in selected regions and was used to detect selection direction. Under a standard neutral model with no change in population size, a strongly negative Tajima’s D value can indicate directional selection removing variation, while a strongly positive value can indicate balancing selection maintaining variation, with 0 reflecting an absence of selection.

For the genetic variation between populations, we used PoPoolation2 v. 1.201 [65] to obtain allele frequency differences for each SNP. The SNPs were then filtered by the following parameters before further analyses: locus coverage > 50 in all populations and an average minor allele frequency (MAF) more than 0.1 [66]. We also obtained pairwise Fst values for non-overlapping 100 kbp windows and for gene sets. We then undertook a principal component analysis (PCA) generated by the prcomp function and package ggbiplot in R [67] based on: 1) MAF across the SNPs after initial filtering as mentioned above; 2) pairwise Fst values from 100 kbp non-overlapping windows to indicate genetic distance patterns across the genome (genome level) [68]; and 3) pairwise Fst for genes (gene level). These Fst estimates were then used to assess patterns of similarity among samples with the same *Wolbachia* infection status in a pairwise comparison, and the same geographic distance in a pairwise comparison.

We further investigated isolation by distance patterns among 2018 samples from the four locations by computing Fst*=Fst/(1 − Fst) [69] based on the averaged pairwise Fst values from 100k bp non-overlapping windows. A geographic distance matrix was built based on the natural log transformation of the shortest road distance between the sampled locations as mosquito movement would be mostly by road transport [70]. We then looked for Fst patterns that might be related to this distance measure and ran a Mantel test through the ade4 package in R to test the relationship between genetic distance and geographic distance [71], with only 2018 populations included.

### Identification of outliers potentially associated with *Wolbachia*

We used two models from a Bayesian outlier approach BayPass v. 2.2 [35] to identify outliers associated with *Wolbachia* infection. Firstly, we used standard covariate model [35, 72], which requires a file providing values of each covariate to produce the Bayes Factor (BF), the ratio of the likelihood of posterior and prior hypotheses, which can quantify the probability of a candidate SNP being under selection [73]. Bayes Factors were exported in dB units (BF* = 10 × log10(BF)); the association with environmental variance was considered “substantial” when BF* > 5, “strongly-selected” when BF* > 10 and “decisively-selected” when BF* > 13 following Jeffreys [73]. We modelled *Wolbachia* infection status as a binary covariate by setting each infected sample as 1 and each uninfected sample as 0. The measures of BFs are based on an Importance Sampling Approximation, which is unstable for single runs in particular when the number of populations is small, so we averaged the BF values of three runs with different seeds for the random number generators following the suggestion in the manual of BayPass [35]. SNPs were considered as “substantial” outliers if the average BF* >= 5 and “strongly-selected” outliers if the average BF* >= 10. The introduction of linear relationships in this model, however, can have high false positives from sampling noise, as the relationships of allele frequency among multiple populations are influenced by genetic drift and migration, and are not statistically independent with each other [36]. Based on this model, we made Manhattan plots for SNPs with average BF* > 0 to show the potential effects of the *Wolbachia* infection at the genome level; SNPs that could not be assigned a position on one of the three autosomes were discarded.

Secondly, we used a BayPass core model to identify outliers from comparisons between GV10 and GV13, GV10 and GV18, given that Gordonvale was the only location where we had samples at time points before and after the release. The infection of *Wolbachia* was considered the main variable across time. An X^T^X algorithm approach was used in this model, which was analogous to an Fst comparison [35, 36]. The X^T^X value was used to identify selection pressure, with higher values under positive selection and smaller values under balancing selection [36]. In this model, we considered SNPs with X^T^X values greater than the 90 % threshold or 95% threshold in both the GV10 and GV13 comparison and the GV10 and GV18 comparison as candidates for intersecting with the “substantial” SNPs and “strongly-selected” SNPs from the first analysis. This model on the other hand, is unable to exclude the noise from gene flow or genetic drift due to the lack of duplicates, which can also cause false positives. We therefore considered the intersection of outliers from the above two models as SNPs potentially associated with *Wolbachia* infection. These SNPs were then matched with GTF annotation file from NCBI to obtain a list of outlier genes.

### Pathway analysis and gene ontology enrichment analysis

We considered SNPs from the intersection of the two BayPass models above as outliers potentially under selection and searched the open reading frame (ORF) to obtain a list of potentially important genes within a 100 kbp region around each SNP (50 kbp upstream and 50 kbp downstream) [74]. These genes were then searched through the KEGG pathway database for interpretation. We then used BLAST search (version 2.9.0 +) in the UniProtKB/Swiss-Prot database [75] through system Entrez to identify homologous proteins in *Drosophila melanogaster*. Only best matches were retained and further filtered with an e-value cut off 1.0E-10 and > 60% identity [76, 77]. GO enrichment analysis was then undertaken in the R packages clusterProfiler [78] and DOSE [79], with a significant false positive rate and false discovery rate cutoff of 0.05.

## Additional files

**Additional file 1**. Genetic and geographic distance matrices. The genetic distance matrix from genome pairwise 100 kbp Fst value, transformed by Fst*=Fst/(1 − Fst), is above the diagonal, the nature log transformed geographic distance (km) is below the diagonal.

|  | GV10 | GV13 | GV18 | EH | RL | YK |
| --- | --- | --- | --- | --- | --- | --- |
| GV10 | 0 | 0.045 | 0.040 | 0.056 | 0.059 | 0.064 |
| GV13 | 0 | 0 | 0.057 | 0.063 | 0.067 | 0.068 |
| GV18 | 0 | 0 | 0 | 0.052 | 0.057 | 0.059 |
| EH | 3.243 | 3.243 | 3.243 | 0 | 0.051 | 0.054 |
| RL | 3.547 | 3.547 | 3.547 | 2.526 | 0 | 0.057 |
| YK | 3.694 | 3.694 | 3.694 | 2.821 | 2.773 | 0 |

**Additional file 2**. IDs of 187 genes where the 950 “substantial” outliers that potentially impacted by *Wolbachia w*Mel infection distributed within.

**Additional file 3**. IDs of 1436 genes within the 100 kbp regions around the 950 outliers that potentially impacted by *Wolbachia w*Mel infection.

**Additional file 4**. Pathways in KEGG database identified from genes potentially impacted by *Wolbachia w*Mel infection

**Additional file 5**. Homologues proteins of *Drosophila melanogaster* in GO enrichment analysis.

**Additional file 6**. Enriched biological process terms.

**Additional file 7**. Enriched cellular component terms.

**Additional file 8**. Enriched molecular function terms.

## Declarations

### Ethics approval and consent to participate

Not applicable.

### Consent for publication

Not applicable.

## Availability of data and materials

The raw data used during the current study belongs to corresponding authors. All data generated during this study are included within this article and its additional files.

## Authors’ contributions

AH and TS conceived the study. ML implemented the study, performed data analysis and wrote the first draft of the manuscript. AH, TS, PAR, QY and LS revised the manuscript. JC and QY provided guidance in data analysis. ML, TS, LS, QY and PAR contributed to sample preparation and/or sequencing. AH secured financial support. All authors read and approved the final manuscript.

## Acknowledgements

We thank Gordana Rasic for her early contribution to the methodology for initial sequencing of the Gordonvale populations. We also thank Moshe Jasper for his advice on data visualization. This research was supported by use of the Nectar Research Cloud, a collaborative Australian research platform supported by the NCRIS-funded Australian Research Data Commons (ARDC).

